# PDZ Domains in Microorganisms: Link Between Stress Response and Protein Synthesis

**DOI:** 10.1101/186510

**Authors:** Vijaykumar Yogesh Muley, Sanjeev Galande

## Abstract

The PSD-95/Dlg-A/ZO-1 (PDZ) domain is highly expanded and diversified in metazoan where it is known to assemble diverse signalling components by virtue of interactions with other proteins in sequence-specific manner. In contrast, in bacteria it monitors protein quality control during stress response. The distribution, functions and origin of PDZ domain-containing proteins in prokaryotes are largely unknown. We analyzed 7,852 PDZ domain-containing proteins in 1,474 prokaryotes and fungi. PDZ domains are abundant in eubacteria; and, this study confirms their occurrence also in archaea and fungi. Of all eubacterial PDZ domain-containing proteins, 89% are predicted to be membrane and periplasmic, explaining the depletion of bacterial domain forms in metazoan. Planctomycetes, myxobacteria and other eubacteria occupying terrestrial and aquatic niches encode more domain copies, which may have contributed towards multi-cellularity and prokaryotic-eukaryotic transition. Over 93% of the 7,852 PDZ-containing proteins classified into 12 families including 6 novel families. Out of these 88% harbour eight different protease domains, suggesting their substrate-specificity is guided by PDZ domains. The genomic context provides tantalizing insight towards the functions associated with PDZ domains and reinforces their involvement in protein synthesis. We propose that the highly variable PDZ domain of the uncharacterized Fe-S oxidoreductase superfamily, exclusively found in gladobacteria and several anaerobes and acetogens, may have preceded all existing PDZ domains.

## Introduction

Proteins exhibiting both signaling and protein interaction domains are prevalent in eukaryotic signal transduction systems (Nourry et al. 2003). This domain architecture provides an elegant solution to rewire and regulate complex biological networks by sensing signals through signaling domains while protein interaction domains serve to amplify the signal. The PDZ domain is one of such protein interaction domains. It was first identified in the context of signaling proteins, which are referred to as GLGF repeats proteins or DHR (Discs large homology repeat) proteins (Cho et al. 1992; Ponting and Phillips 1995). The abbreviation PDZ is derived from the three metazoan proteins in which this domain was first identified: **P**SD-95, **D**LG and **Z**O-1(Kennedy 1995).

Metazoan PDZ domains are referred to as canonical PDZ domains and typically comprise 80-100 amino acid residues harboring a highly conserved fold (Doyle et al. 1996; Kennedy 1995). However, the length of secondary structures composed of six β-strands with a short and a long α-helix may vary (Doyle et al. 1996; Morais Cabral et al. 1996). In contrast, eubacterial PDZ domains fold similarly to metazoan domains but with a distinct topology of secondary structural elements and are referred to as non-canonical (Harris and Lim 2001; Lee and Zheng 2010). Non-canonical PDZ domains consist of a circularly permuted structural fold.

PDZ domains are inherently variable at primary sequence level, and show diversity in functional roles and binding specificities in metazoan (Belotti et al. 2013; Chen et al. 2008; Sakarya et al. 2007; Sakarya et al. 2010). The origin, diversity and functions of prokaryotic and fungal PDZ domains are largely unknown. The genome-wide analysis of non-metazoan PDZ domains dates back to 1997, wherein these domains were shown to occur in bacteria (in abundance), plants and fungi [13]. However, its presence in fungi was considered doubtful due to low sequence similarity with known PDZ domains. Therefore, it was assumed either that the primordial PDZ domain arose prior to the divergence of bacteria or eukaryotes, or that horizontal gene transfer led to the acquisition of these domains by bacteria [13]. Even until now few researchers believe that this domain is occurs in fungi and archaea [12], while few others do not [7, 14], indicating how little we know about non-metazoan PDZ domains. It was hypothesized that a subset of a eubacterial PDZ domains might form precise canonical structures observed in metazoans, and the domain might have co-evolved with multi-cellularity and organismal complexity (Harris and Lim 2001; Sakarya et al. 2007). However, the PDZ domain-containing protein repertoire in prokaryotes and fungi is not explored at genomic levels to clarify such contradictions and hypotheses.

In this study, we have identified and analyzed an entire set of PDZ domains in 1,474 prokaryotic and fungal genomes using cutting-edge remote homology detection sequence analysis tools. Several eubacteria code for more than fifteen PDZ domain-coding genes, and have complex structures unusual to bacteria and certain forms of multicellular communities. We classified 97% of these proteins into 12 families based on conserved domain architecture, of which eight have been reported here for the first time, providing a glimpse of ancestral history and functional clues. Genomic context analysis connects these genes to protein synthesis. This work bridges the ever-increasing gap between prokaryotic/fungal, and metazoan PDZ domain studies.

## Results

### PDZ domains are ubiquitous and occur even in archaea and fungi

The 7,852 proteins containing altogether 9,975 PDZ-like domains were identified in 1,419 of 1,474 (96%) genomes. We observed an abundance of domain copies in eubacteria and archaea but mostly single incidence in fungal genomes (Fig. 1 & Table 1). Eubacterial species encode significantly higher numbers of domains than archaea and fungi with Wilcox test p-value 2.2e-16 and 2.769e-06, respectively (Fig. 2a). Our results are mostly consistent with previous findings; however, we contradict the domain’s absence in archaea and fungi (Harris and Lim 2001; Ponting 1997).

**Figure 1.**
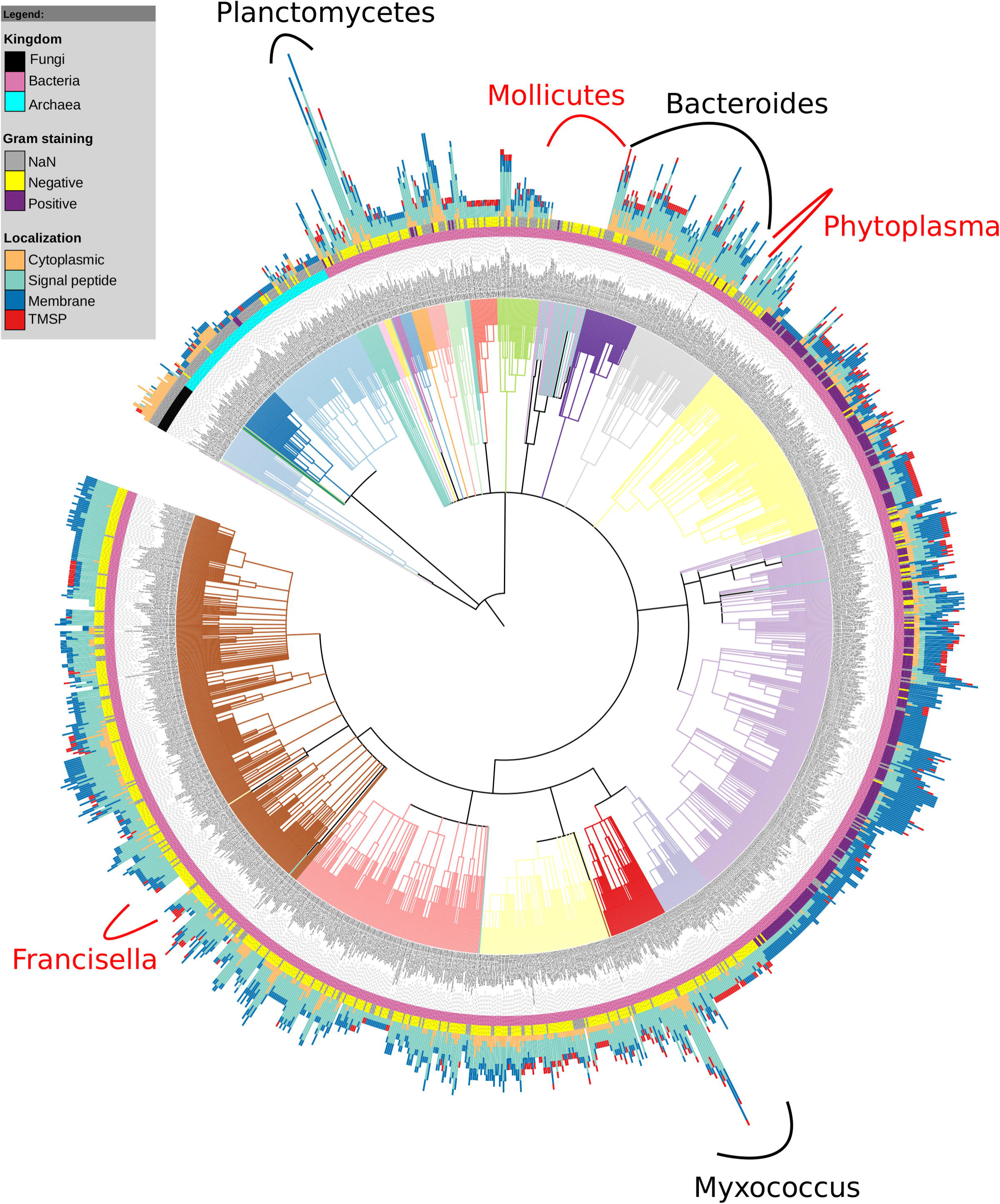
The phylogenetic extent of PDZ domain-containing proteins across prokaryotes and fungi. The distribution of PDZ domain-containing proteins is superimposed over the NCBI taxonomy tree using interactive Tree of Life (iTOL) web server. The outermost layer depicts bar plots of a height proportional to the number of PDZ domain-containing proteins in a taxon. Each bar plot features four colors corresponding to the number of proteins in their four predicted sub-cellular locations. The innermost and middle color strips represent Gram nature and kingdom of taxa, respectively. Leaves are colored to distinguish various classes/phyla. Lineages encoding a higher number and no PDZ domains are annotated in black and red colors, respectively. The distribution shows ubiquitous occurrence of PDZ proteins across bacteria, and scarcity in archaea and fungi. A higher number of membrane and secretory proteins are present in Gram-positive and-negative bacteria, respectively. Fungi encode mostly cytoplasmic PDZ proteins.

**Figure 2.**
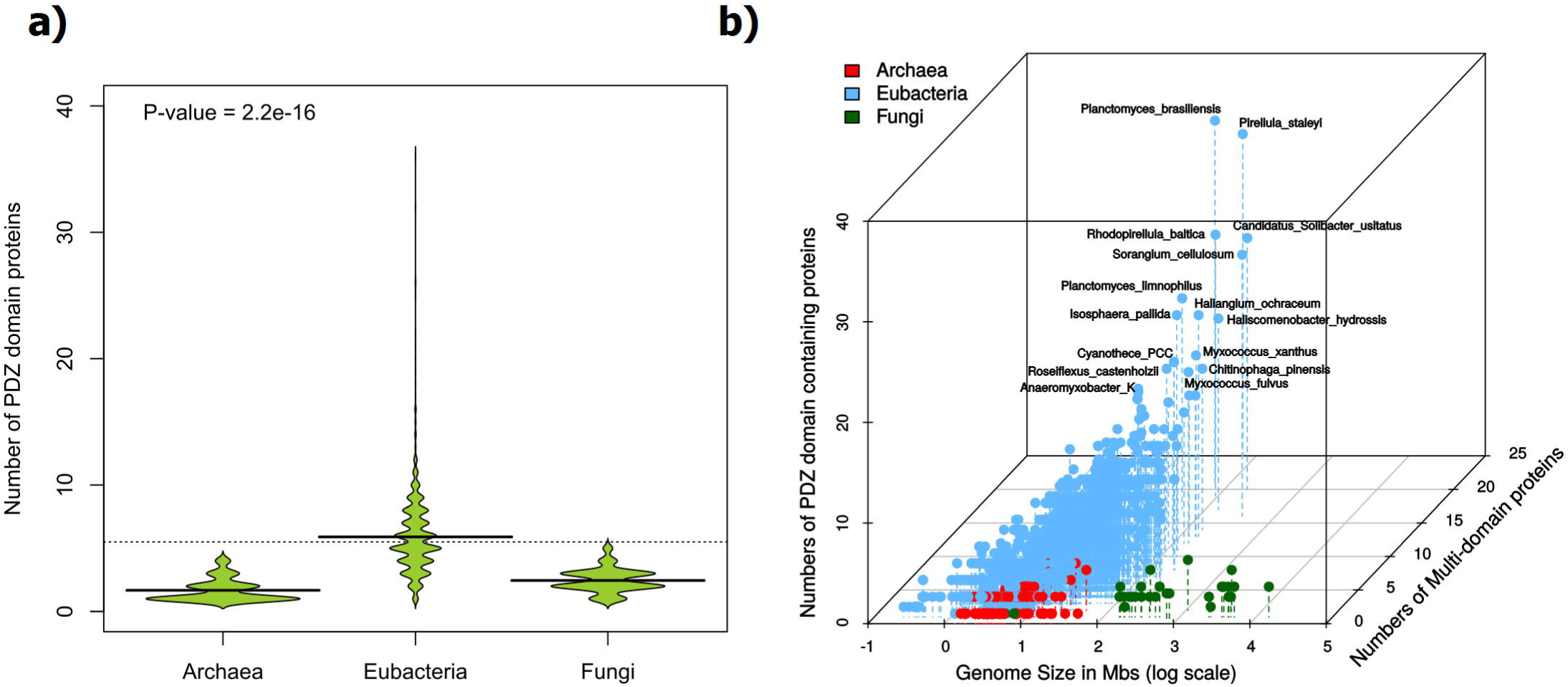
The occurrence of PDZ domain-containing proteins and its relationship with genome size in eubacteria, archaea and fungi. a) The bean plot demonstrates the distribution of proteins in the three domains. In the plot, the horizontal dotted line corresponds to the overall average number of proteins across all genomes under study. Bean-width corresponds to the proportion of genomes in it and bean-line shows the average number of proteins in the kingdom. Archaeal and fungal genomes encode less than 5 proteins in average, whereas many eubacteria encode a higher number of proteins. b) The scatter plot illustrates the relationship between genome size, number of proteins and multi-domain proteins. Numbers of proteins and their multi-domain architecture expanded with increase in genome size in eubacteria. Proteins with repetitive PDZ domain alone were not counted as multi-domain. The species with complex processes and/or the ability to form cell aggregates are annotated in the plot.

**Table 1.** Taxonomic distribution of PDZ domain-containing protein families classified in this work.

| Kingdom | Phylum | Htr | Ctp | RseP | APN<sup>§</sup> | Lon<sup>§</sup> | GspC | FeSo<sup>§</sup> | SpoIVB | AP<sup>§</sup> | ComP | ChaN<sup>§</sup> | ZEP<sup>§</sup> |
| --- | --- | --- | --- | --- | --- | --- | --- | --- | --- | --- | --- | --- | --- |
| Fungi |  | 28/25 | - | 2/2* | - | - | - | - | - | - | - | - | - |
|  | Ascomycota | 28/25 | - | - | - | - | - | - | - | - | - | - | - |
|  | Basidiomycota | - | - | 2/2* | - | - | - | - | - | - | - | - | - |
|  | Microsporidia | - | - | - | - | - | - | - | - | - | - | - | - |
| Archaea |  | 55/43 | 5/5* | 109/108 | 13/13 | - | - | - | - | - | - | - | - |
|  | Euryarchaeota | 27/18 | 2/2* | 75/74 | - | - | - | - | - | - | - | - | - |
|  | Crenarchaeota | 27/24 | 3/3* | 33/33 | 13/13 | - | - | - | - | - | - | - | - |
|  | Nanoarchaeota | - | - | - | - | - | - | - | - | - | - | - | - |
|  | Other Archaea | 1/1 | - | 1/1 | - | - | - | - | - | - | - | - | - |
| Eubacteria |  | 2902/1271 | 1716/1041 | 1262/1242 | 294/246 | 292/290 | 182/170 | 156/148 | 139/136 | 49/48 | 48/41 | 37/37 | 29/25 |
|  | Acidobacteria | 20/6 | 23/6 | 6/6 | 10/5 | - | - | - | - | 1/1 | 8/6 | - | 2/2 |
|  | Actinobacteria | 302/138 | 31/24 | 117/116 | 1/1 | 113/113 | - | 5/5 | - | - | - | - | - |
|  | Alphaproteobacteria | 470/146 | 165/128 | 146/145 | 29/19 | - | 11/11 | - | - | 2/2 | - | 1/1 | - |
|  | Aquificae | 13/9 | 11/9 | 10/9 | - | - | 3/3 | - | - | - | - | 1/1 | - |
|  | Bacteroidetes/Chlorobi | 81/63 | 265/63 | 63/63 | 33/22 | - | - | - | - | 22/22 | 1/1 | - | 7/7 |
|  | Betaproteobacteria | 297/96 | 116/96 | 94/94 | 84/84 | - | - | - | - | 6/6 | - | 7/7 | 4/4 |
|  | Chlamydiae/Verrucomicrobia | 21/20 | 24/20 | 20/20 | - | - | - | - | - | - | - | - | - |
|  | Chloroflexi | 45/15 | 38/15 | 15/15 | 6/6 | - | - | 7/7 | - | - | 4/4 | - | - |
|  | Cyanobacteria | 121/40 | 112/40 | 41/40 | 29/29 | - | - | 40/40 | - | - | - | - | - |
|  | Deinococcus-Thermus | 47/13 | 30/13 | 13/13 | - | - | - | 6/6 | - | - | - | - | - |
|  | Deltaproteobacteria | 120/43 | 77/43 | 52/43 | 10/7 | - | 23/21 | 9/9 | - | 5/5 | - | 17/17 | 4/4 |
|  | Epsilonproteobacteria | 38/37 | 38/38 | 38/38 | - | - | 1/1 | - | - | - | - | - | - |
|  | Firmicutes | 446/290 | 258/200 | 287/287 | - | 178/176 | - | 84/76 | 137/134 | - | 34/29 | - | - |
|  | Fusobacteria | 4/2 | 5/5 | 4/4 | - | - | - | - | - | - | - | - | - |
|  | Gammaproteobacteria | 663/282 | 416/270 | 282/275 | 85/66 | 1/1 | 137/127 | - | - | 10/10 | 1/1 | 8/8 | 3/3 |
|  | Other Bacteria | 65/23 | 39/25 | 26/26 | 1/1 | - | 1/1 | 5/5 | 2/2 | 1/1 | - | 3/3 | - |
|  | Planctomycetes | 55/5 | 19/5 | 5/5 | - | - | - | - | - | 2/1 | - | - | 9/5 |
|  | Spirochaetes | 82/31 | 37/29 | 31/31 | 6/6 | - | 6/6 | - | - | - | - | - | - |
|  | Thermotogae | 12/12 | 12/12 | 12/12 | - | - | - | - | - | - | - | - | - |

A protein family is shown by its total number of members/total number of organisms in particular taxonomic group. Protein families are abbreviated as : Htr, High temperature requirement proteases; Ctp, C-terminal processing proteases; RseP, RseP intramembrane metalloproteases; APN, Aminopeptidase protein N; Lon, ATP-dependant lon proteases; SpoIVB, Sporulation factor IV B proteases; AP, Aspartate proteases; ZEP, Zn-dependent exopeptidases; GspC, General secretion pathway protein C; FeSO, Fe-S oxidoreductases; ComP, Competence protein family; ChaN, EreA-Chan-like family.
<sup>§</sup> Uncharacterized families.
\* Rare incidences of family proteins in respective group/kingdom which were considered as horizontally transferred events from eubacteria.

Many PDZ domain-containing proteins are reported to be periplasmic or intra-membrane (Dartigalongue et al. 2001; Dong and Cutting 2004; Lipinska et al. 1990). Interestingly, the domain is absent in 4% species (55 genomes), mainly belonging to cell-wall lacking mollicute and candidatus phytoplasma species, of which many are adapted to parasitic lifestyle (Fig. 1). The domain is also absent in two *Buchnera aphidicola* species which are symbionts of a Cinara (a conifer aphid) and in the archaean *Nanoarchaeum equitans*, a parasite on *Ignicoccus hospitalis* on which it depends for lipids (Jahn et al. 2008). Buchnera species lost genes encoding cell-surface components such as lipopolysaccharide (LPS) and phospholipids which are present in its close relatives (Shigenobu et al. 2000). These observations suggest that the lack of cell envelope components may be related to the absence of the domain. Thus, we studied the predicted sub-cellular localization of the domain-containing proteins. Interestingly, 88.84% proteins are predicted to have signal peptides, transmembrane helices or both, whereas the remaining 11.16% were cytoplasmic (Supplementary Fig. S1a). The presence of a signal peptide indicates secretion of the protein or insertion into membrane compartments. These results associate eubacterial proteins mainly to the membranes and membrane compartments (Supplementary Fig. S1b), and could explain their absence in membrane lacking species. The depletion in archaea and fungi could be due to their different membrane architecture (Supplementary Fig. S1b) (Lombard et al. 2012). In these organisms, the lack or change in cell envelope components might have relieved functional constraints on non-canonical PDZ domain-encoding genes with their subsequent loss during evolution.

Sub-cellular localization of proteins differs between Gram-negative and-positive bacteria since the later do not possess a double membrane with periplasmic space between them. A significantly higher number of proteins with a signal peptide was observed in Gram-negative species whereas transmembrane helices occurred more frequently in Gram-positive bacteria, with Wilcox test p-values 2.2e-16 (Supplementary Fig. S1c). Gram-positive species also have significantly lower numbers of proteins compared to Gram-negative (Wilcox test p-value 1.925e-08), suggesting either they have lost the additional PDZ proteins when their ancestors lost one of the two membranes during evolution (Cavalier-Smith 2006) or never had them. In former case, the ancestral cells which incorporated the PDZ domain-containing proteins into a single membrane might have had selective advantages. Collectively, our results indicate wide occurrence of PDZ domains in prokaryotes and fungi. Their absence in certain species could be evolutionarily linked to the loss of cell envelope components.

### Cellular complexity, multi-cellularity and abundance of PDZ proteins

Eubacterial PDZ proteins are primarily involved in stress signaling. The number of signaling proteins in prokaryotes is proportional to their genome size (Galperin 2005). PDZ domains also follow this trend. Figure 2b shows that numbers of PDZ domain-containing proteins and their multi-domain architecture expanded with increase in genome size. However, this trend is significant only for eubacterial genomes and not for archaea and fungi (Supplementary Fig. S2).

PDZ being the second most abundant domain in metazoa, it was hypothesized that the domain might have co-evolved with multi-cellularity and complexity (Harris and Lim 2001; Kim et al. 2012). Interestingly, many eubacterial species encode more than 10 PDZ genes, particularly planctomycetes and myxobacteria (Fig. 1 & 2b). Interestingly, besides methylotrophic proteobacteria, only planctomycetes and myxobacterial members can synthesize C_30_ sterols such as lanosterol, which are found primarily in eukaryotes (Desmond and Gribaldo 2009; Fuerst and Sagulenko 2011; Pearson et al. 2003). Planctomycetes are also considered a probable host for an endosymbiont which gave rise to a eukaryotic cenancestor cell, as they exhibit many features found only in eukaryotes such as internal membranes, a primitive form of endocytosis, growth by budding and lack of peptidoglycans (Fuerst and Sagulenko 2011; Godde 2012). Myxobacteria are known to form fruiting bodies which behave in many aspects like a multi-cellular organism (Rokas 2008), and thus may demarcate the origin of multi-cellularity. Ser/Thr/Tyr protein kinases, which together comprise a major class of regulatory proteins in eukaryotes, were ubiquitously found in myxobacteria and planctomycetes (Perez et al. 2008). Corroborating our results with above-mentioned findings raises the question, whether PDZ domain expansion in planctomycetes and myxobacteria to metazoan extent is an evolutionary contribution towards the origin of eukaryotes and multi-cellularity. We speculate the possible contribution of planctomycetes and myxobacteria to the evolution of a eukaryotic cenancestor cell, which diverged into fungi that lost PDZ domains, and a probable metazoan progenitor that retained PDZ domains and expanded subsequently during metazoan evolution. Nonetheless, our results strengthen the hypothesis that the expansion of PDZ domains indeed correlates with multi-cellularity and complexity - even in eubacteria.

Metazoan life flourished in limited ecological niches compared to the extremely diverse habitats of prokaryotes. Remarkably, prokaryotes that are most successful in terrestrial and aquatic ecological niches encode significantly higher numbers of PDZ domains compared to those inhabiting multiple environments; those that live obligatorily host-associated; and those with specialized habitat, i.e. environments such as marine thermal vents (Fig. 3a). Evidently, prokaryotes encoding higher number of PDZ domains are aerobic rather than anaerobic or facultative with Wilcox p-value 2.1e-05 (Fig. 3b). These results indicate that the expansion of PDZ domain coding genes might have role in adapting to aerobic aquatic and terrestrial habitats.

**Figure 3.**
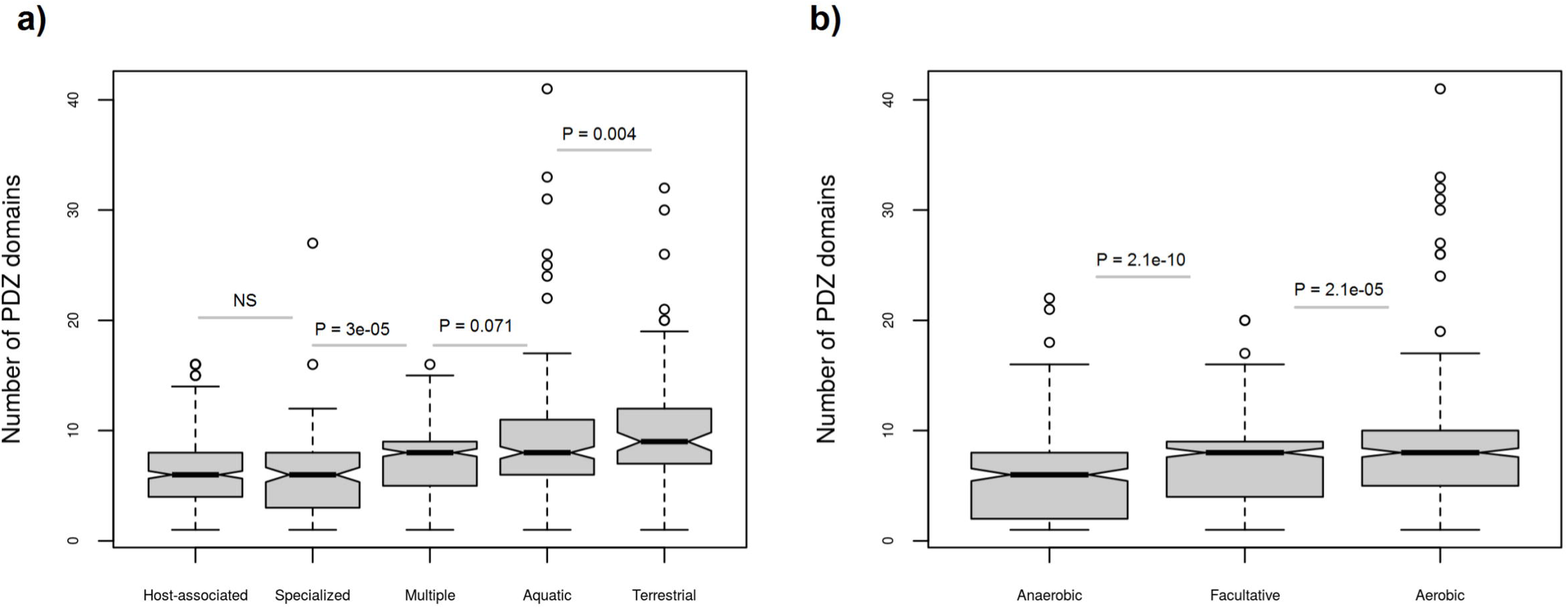
Eubacterial species encoding higher numbers of PDZ domains occupy habitats similar to metazoan. The boxplot depicts the distribution of PDZ domains in prokaryotes that are classified based on their (a) habitats and (b) oxygen requirement. Test of significance p-values using Wilcox test are shown above of compared two groups; null hypothesis was the number of PDZ domains in organisms belonging to right side group is higher than the left side. NS stands for non-significant p-value. Prokaryotes encoding a higher number of PDZ domains favor aquatic, terrestrial and aerobic niche. In panel b) “Multiple” stands for species with a wide host range and variety of habitats; “Host-associated” for species obligatorily associated with a host; and “Specialized” for those with specialized habitat, i.e. environments such as marine thermal vents.

### Domain architecture and classification of PDZ proteins

Superfamily database classified PDZ domain-like sequences into six families based on subtle differences in their protein structures, which are: *PDZ domain*, the metazoan canonical form; *High Temperature Requirement A-like serine protease (HtrA); Tail specific protease (Tsp); EpsC C-terminal domain-like; MTH1368 C-terminal domain-like; and Interleukin 16 domain-like* (Supplementary Fig. S3a) (Gough et al. 2001). Likewise, Pfam database grouped domain sequences based on sequence similarity into three families: *PDZ*, the metazoan canonical form; *PDZ_2*, found in plants and eubacteria; and *PDZ_1*, found in few members of fungi and archaea (Supplementary Fig. S3b) (Sonnhammer et al. 1997). We confirm widespread occurrence of the non-canonical HtrA superfamily and PDZ_2 Pfam domains across eubacteria (Ponting 1997), as well as an abundance of the canonical PDZ domains as hypothesized by Harris and Lim, which might have contributed towards expansion and diversity in metazoan counterparts (Supplementary Fig. S3) (Harris and Lim 2001).

We inspected similar domain architectures in semi-automated fashion (detailed in ‘Methods’) and classified 7,318 proteins out of 7,852 into twelve families, constituting 97% of all (Fig. 4a, please see Supplementary Dataset 1 for individual proteins’ details). The remaining proteins, represented in less than 20 species were treated unclassified. The PDZ domain plugged at either the N-or C-terminus of other domains with no apparent preference for one of the termini (Fig. 4a). Out of twelve classified protein families, PDZ domain is combined with eight different protease domains in 88% of the classified proteins. The remaining proteins classified into four non-protease families and referred to as Fe-S oxidoreductase, General Secretory Protein C, Haem-binding uptake, and Sensor histidine kinase ComP, based on the functions of the domains therein (Fig. 4b). Figure 4c shows the distribution of proteins into eight protease families suggesting that the HtrA, Carboxy-terminal protease (Ctp) and Regulator of Sigma-E Protease (RseP) families proteins are predominant compared to lesser number of proteins belonging to Aminopeptidase N, Lon Protease, Sporulation protein IV B, Aspartate protease, and Zn-dependent exopetidases families. In spite of sequence and structural diversity, PDZ domain is preferentially combined with protease domain in 88% of classified proteins, suggesting its function to be directly related to providing substrates for proteolysis. Our assumption is supported by the function of PDZ domains of well-characterized proteases HtrA, Ctp, and RseP in *Escherichia coli* (Anbudurai et al. 1994; Clausen et al. 2011; Li et al. 2009).

**Figure 4.**
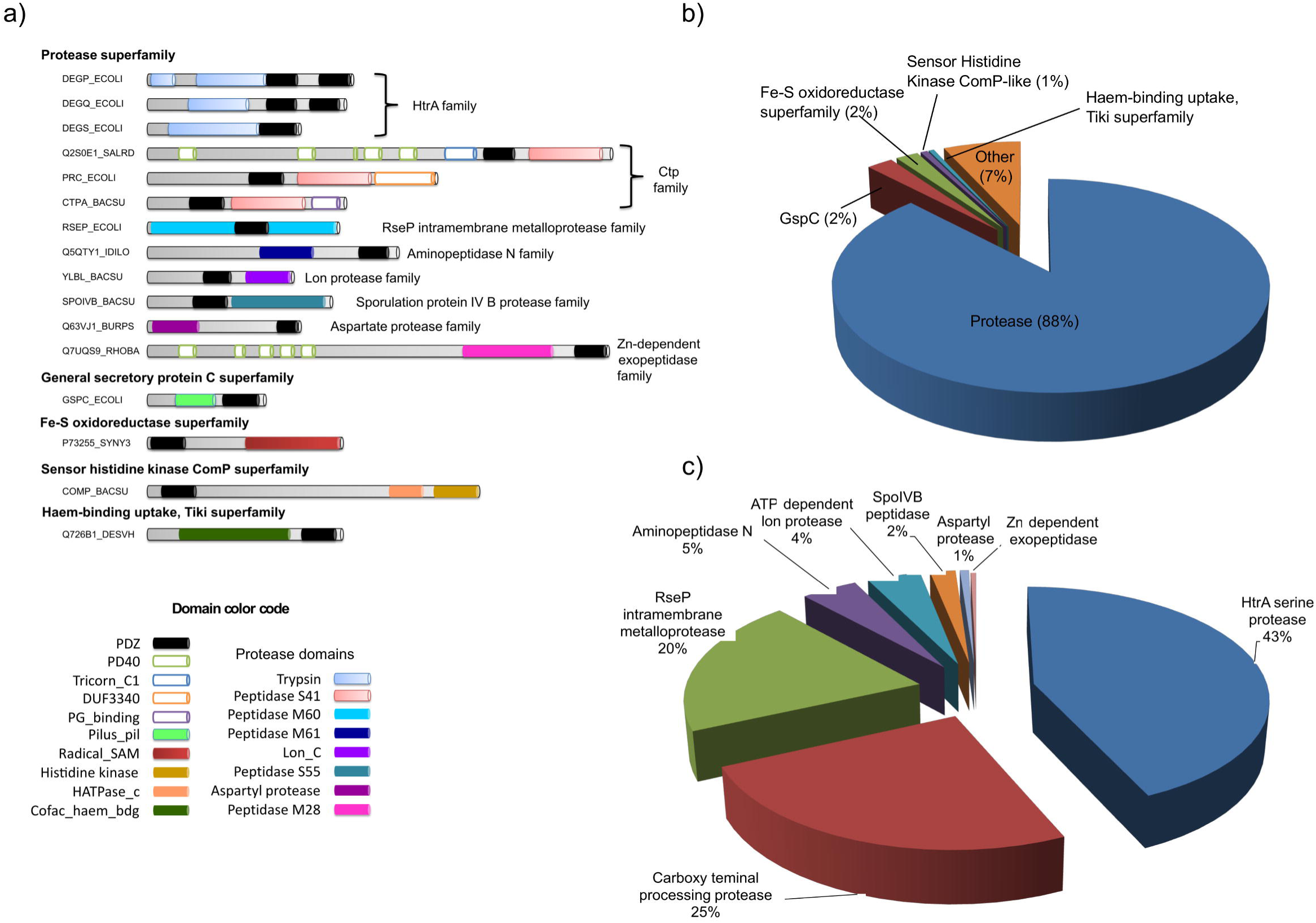
Modular organization and classification of PDZ domain-containing proteins. a) Panel depicts the most frequently observed domain architectures of PDZ domain-containing proteins. Proteins are scaled approximately to the length of their primary sequence. Each domain architecture is a representative candidate of the superfamily/family/sub-family identified in this work. Protein identifiers are UniProt accession names. The Trypsin domain of the DEGP_ECOLI protein is interrupted by a Q-linker stretch. Protease domains are frequently combined with PDZ. b) The distribution of all PDZ domain-containing proteins into super-families identified in this study based on common domain architectures shown in (a). c) The protease superfamily members are grouped into families based on protease type. Domain abbreviations: PD40 for WD40-like Beta Propeller Repeat domain; Tr_C1 for Tricorn protease C1 domain; Tr_PDZ for Tricorn protease PDZ domain; PS41 for Peptidase family S41; DUF3340 for C-terminal domain of tail specific protease; PG_b for Peptidoglycan binding 1 domain; PM50 for Peptidase family M50; PM61 for Peptidase family M61; Lon_C for lon protease C-terminal proteolytic domain; PS55 for Peptidase family S55; Asp_prt for Aspartyl protease domain; PM28 for Peptidase family M28; Pilus_P for Type IV pilus biogenesis domain; Radical_SAM for Radical SAM superfamily; HisKA_3 for Histidine kinase; HATPase_c for Histidine kinase; DUF399 for Domain of unknown function.

The phyletic distribution of classified proteins is shown in Table 1. All families are represented in eubacteria; three also occur in archaea; and the HtrA family is common even in fungi. Notably, HtrA and Ctp family proteins are present in multiple copies in eubacteria suggesting their expansion by gene duplication during evolution (Table 1). Notably, fungi inherited HtrA genes only, which encode non-canonical PDZ domains. It is consistent with their presence in metazoans and inheritance from last common ancestors of eukaryotes. Some of the interesting families have been discussed in detail in the subsequent sections.

### High Temperature Requirement A (HtrA)-like serine proteases: presence in the three domains of life

The HtrA protein family is part of the elaborate high temperature stress response system for protein quality control, which monitors protein homeostasis to prevent accumulation of unwanted and damaged proteins in the cell (Clausen et al. 2011). Characteristic feature of HtrA proteins is one or two PDZ(s) followed by a trypsin-like serine protease domain (Fig. 4a). Previous studies have reported its presence in higher eukaryotes (Pallen and Wren 1997). HtrA-like proteins are present in 90% of the analyzed prokaryotic and fungal genomes, confirming their presence in the three domains of life. The expansion of HtrA family proteins in most eubacteria phyla might have been selected through membrane stress, especially in high temperature (Table 1).

### Carboxy-terminal protease (Ctp) family: bacterial root of metazoan PDZ domains

The characteristic feature of Ctp family proteins is the presence of canonical or non-canonical PDZ domain followed by a Peptidase_S41 domain. The PDZ domain of a Ctp recognizes four hydrophobic residues at the C-terminus of the FtsI precursor, where Peptidase_S41 cleaves the C-terminus and produce mature FtsI (Hara et al. 1991) (Fig. 4a). The PDZ and Peptidase_S41 domain combination is observed in 1,721 proteins accompanied by many other domains in lineage-specific manner. These are Domain of Unknown Function 3340 (DUF3340), Peptidoglycan binding domain, Tricorn and Tricorn_C1 domains (Fig. 4a). This suggests that Ctp family is amenable to evolutionary changes more than others, and associated domains provided new functional aspects to it in lineage-specific manner.

The canonical PDZ domain was detected based on Pfam as well as Superfamily profile in 309 (out of 316) DUF3340 domain-containing family proteins in 298 species (Supplementary Table S1). Interestingly, the Ctp DUF3340 domain shows sequence similarity with the mammalian interphotoreceptor retinoid binding protein (IRBP) (Keiler and Sauer 1995) and might be ancestral to it (Ponting 1997). We searched the DUF3340 region of *Escherichia coli* Ctp (Prc) in UniProt database and found similar sequences in five archaeal proteins and many metazoan species including the early branch point’s demosponge, sea anemone, and also in human. However, we were unable to detect similar sequences in fungi and ecdysozoa members-worm and fly. This suggests that the possible transfer of eubacterial PDZ domains in metazoan along with DUF3340 (IRBP). Archaea presumably never had canonical PDZ domains. The bacterial endosymbiont might have contributed these domains to eukaryotic phylogeny, where mainly they were lost recurrently just in fungi and ecdysozoa. Moreover, phylogenetic analysis discussed in subsequent section confirms recent divergence of the PDZ domains of Ctp family.

### Uncharacterized members of Fe-S oxidoreductase superfamily: at the base of PDZ domain evolution

A highly variable form of the PDZ domain, which is beyond recognition by Pfam sequence profile but predicted with the structure-based Superfamily HMM profile, is observed along with Pilus_pil and Radical SAM domains in several proteins. The Pilus_Pil domain containing proteins was found in 170 Gram-negative eubacterial species, mostly in proteobacteria. The prototype example is General Secretory Protein C (GspC) of *Escherichia coli*, a part of type II secretion machinery (Francetic et al. 2000). On the other hand, Radical SAM domain observed in 156 uncharacterized proteins in 148 eubacterial species. These are referred to as Fe-S oxidoreductases due to the highly conserved iron-sulphur binding motif and radical SAM domain. One PDZ domain occurs at the N-terminus of Radical SAM domain containing uncharacterized Fe-S oxidoreductase proteins with a CxxxCxxC conserved motif in 148 species (Supplementary Fig. 4). The amino acids responsible for structural integrity seem to be conserved in these two families. It could be a result of accumulated mutations over time, and hence detected by the structure-based HMM model of PDZ domains only. Thus, we hypothesized that one of the PDZ domain of these two families could be ancestral to all other PDZ domains.

GspC family is a part of general secretory system, which transports many substrates. This system is not specific towards certain substrate hence naturally imparts less constraints on the sequence conservation. GspC family proteins are mostly present in proteobacteria, which is not accounted among old bacterial phyla so far. On the other hand, the radical SAM domain of Fe-S oxidoreductase family is considered to be among the oldest domains. The highly conserved CxxxCxxC motif is likely to form an iron-sulphur (Fe-S) cluster to cleave SAM reductively and produce a radical (usually a 5′-deoxyadenosyl 5′-radical) (Frey et al. 2008). The radical intermediate allows wide variety of unusual chemical transformations. These proteins are conserved in all cyanobacterial genomes investigated in this study, and half of the chlorobacteria and hadobacteria (Table 1). These phyla constitute the oldest bacterial clade gladobacteria (Cavalier-Smith 2006) and a probable root of life is placed in chlorobacteria (Cavalier-Smith 2006). These proteins are also present in firmicutes, δ-proteobacteria; and few actinobacterial species, which predominantly use anaerobic mode of respiration (Table 1). The presence of this family’s proteins in photosynthetic and anaerobic eubacteria suggests their functional role in the early Earth environment (Lane and Martin 2012) and also is consistent with their high sequence divergence due to long evolutionary time span. Furthermore, it is presents in several acetogens that generate acetate as a product of their anaerobic respiration and their modern lifestyles resemble to that of last universal common ancestor (Weiss et al. 2016). MSA analysis confirms the authenticity of these PDZ domains following structural analysis (Fig. 5a). We used protein sequence of slr2030 and GSU1997, hypothetical proteins from *Synechocystis* PCC 6803 and *Geobacter sulfurreducens* PCA to predict three-dimensional structure using Phyre2 protein modelling web server. The 61% N-terminus amino acids of both sequences could be modelled with more than 90% confidence level, which includes PDZ and radical SAM domain. The predicted structures were compared with the known ligand-bound PDZ domain structure of HtrA2 protein isolated from *Mycobacterium tuberculosis* (Mohamedmohaideen et al. 2008) (Fig. 5b). The predicted domain structure is composed of two helices and three to four beta sheets as opposed to around six often occur in known structures (Fig. 5c, d). This indicates the insertion of additional beta sheets in PDZ domains of other families later in the evolution. Even with only three beta sheets the peptide-binding cavity in the domain is maintained suggesting its functional equivalence to other PDZ domains (Fig. 5c, d). The overlap between predicted PDZ structures of slr2030 and GSU1997 proteins with the HtrA2 (PDB id 2z9i) is very high with root mean square deviation of 0.35 and 0.61, respectively. We performed phylogenetic analysis to infer the divergence and origin of the domain. The PDZ domains of Fe-S Oxidoreductases consistently placed at the root by four different phylogeny reconstruction methods, while Ctp PDZ domain at the tip suggesting its recent divergence (Fig. 5e, Supplementary Fig. 5,6,7). On the other hand, GspC family domains may have diverged later from HtrA family domains.

**Figure 5.**
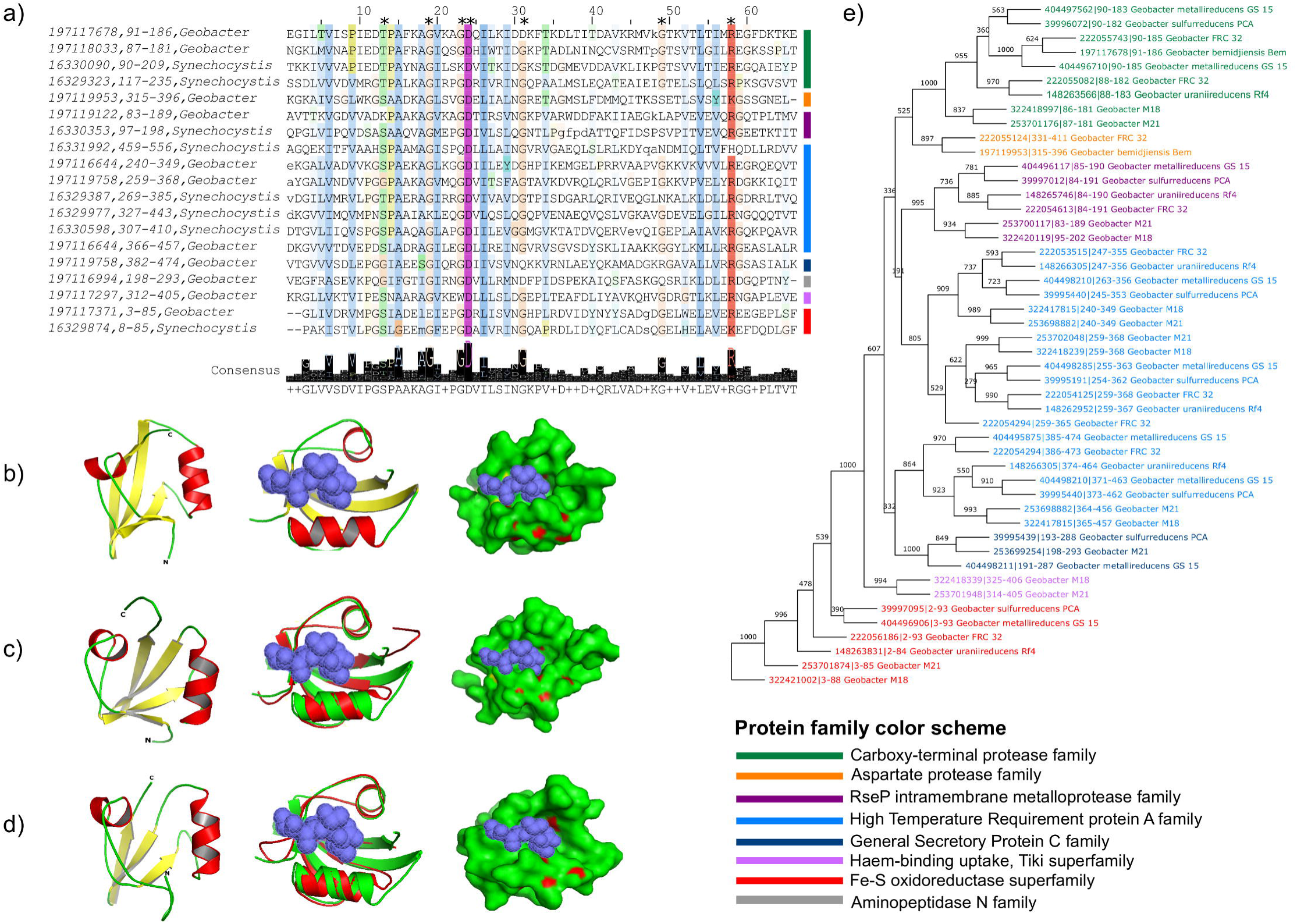
Structure, function and evolution of uncharacterized Fe-S oxidoreductase superfamily. a) Multiple sequence alignment of PDZ domains from Geobacter and Synechocystis. Positions with asterisk marks on top in MSA are also conserved in metazoan PDZ domains. b) The structure of PDZ domain of HtrA2 protein from *Mycobacteria* is shown without and with tetra-peptide ligand in cartoon, and spacefill model. The predicted structure of PDZ domain from Geobacter (c) and Synechocystis (d). Their overlap with HtrA2 is shown along with its ligand. HtrA2 is shown in green color while predicted structure in red. The predicted structure bound with HtrA2 ligand is shown in space-fill model in third panel. Phylogenetic tree shown in panel (e) is constructed by Fitch neighbor-joining algorithm. Phylogenetic tree shows ancestry of PDZ domain of Fe-S oxidoreductases to other PDZ domains, whereas Ctp family domain as a recent divergence.

Collectively, phylogenetic analysis strongly indicates that the PDZ domain of radical SAM proteins is likely to be ancestral and gave rise to other domains, of which Ctp family domain is diverged recently. Phyletic distribution also supports the presence of PDZ domains of radical SAM proteins in the last universal common ancestor.

### Genome neighbourhood analysis: connection with protein synthesis and membrane remodelling

Conserved chromosomal co-localization of genes reflects their co-regulation and often their products participate in the same functions. The genomic neighbourhood of genes encoding the PDZ domain-containing Regulator of Sigma-E Protease (RseP), ATP-dependent lon proteases (in *Bacillus subtilis* known as YlbL) and radical SAM family is conserved in phylogenetically diverse species (Fig. 6). RseP is an inner membrane metalloprotease induces stress response via σ24 (RpoE) factor upon its activation by HtrA family protein DegS in response to damaged outer membrane proteins in *Escherichia coli* (Dartigalongue et al. 2001; Li et al. 2009). RseP-like proteins are highly conserved (92% genomes) among other PDZ domain-containing family proteins (Table 1). Characteristic feature of this family’s proteins are two highly conserved motifs, HExGH and NxxPxxxLDG at their N-and C-termini which are similar to the potential zinc binding site found in a variety of metalloproteases (Brown et al. 2000) and to the motif found in the human S2P protease, respectively (Supplementary Fig. S8) (Dartigalongue et al. 2001). The proximal genes of RseP mostly encode proteins responsible for outer membrane biogenesis and translation. These include bamA (OMP biogenesis) (Rhodius et al. 2006), cdsA (lipid biosynthetic pathway), uppS (or ispU, essential for carrier lipid formation in bacterial cell wall synthesis) (Kato et al. 1999), dxr (upstream of uppS catalyzed pathway), rpsB (part of the 30S ribosomal subunit), and frr (ribosome recycling factor upon translation termination) (Hirokawa et al. 2004) (Fig. 6a). Supplementary Figure S9 depicts the complex regulation of this gene cluster in *Escherichia coli* by many σ24 (a stress response sigma factor) promoters, (Erickson and Gross 1989); σ70, a primary sigma factor during exponential cell growth; and also rho-independent termination sites. The presence of the frr gene is especially interesting since some of its mutants rapidly decrease protein synthesis followed by inhibition of RNA synthesis at 42 ^o^C (Hirokawa et al. 2004). Mutations in rseP characteristic motifs also have been shown to be lethal at temperatures above 41 ^o^C (Dartigalongue et al. 2001). The upstream region of both rseP and frr genes has a σ24 promoter, suggesting their co-regulation (Supplementary Fig. S9). Therefore, we propose a possible link between translation and membrane biogenesis regulated by the stress response factor σ24, mediated by the induction of rseP gene expression along with HtrA family member’s degP and degQ during heat shock. σ24 is positively regulated by the starvation alarmone ppGpp (guanosine 3′-diphosphate 5′-diphosphate) during entry into stationary phase, suggesting that σ24 can respond to internal signals as well as stress signals originating in the cell envelope (Costanzo and Ades 2006). We propose the internal signal through ppGpp might be responsible for the σ24 activation to maintain misfolded and damaged proteins during stationary phase (internal signal) and for inhibition of *tff-rpseB-tsf* gene expression to cease translation due to nutrient limitations, since ppGpp has been shown to inhibit expression of this transcription unit during amino acid starvation (Aseev et al. 2014). These observations associate RseP family proteins directly to translation and membrane biogenesis.

**Figure 6.**
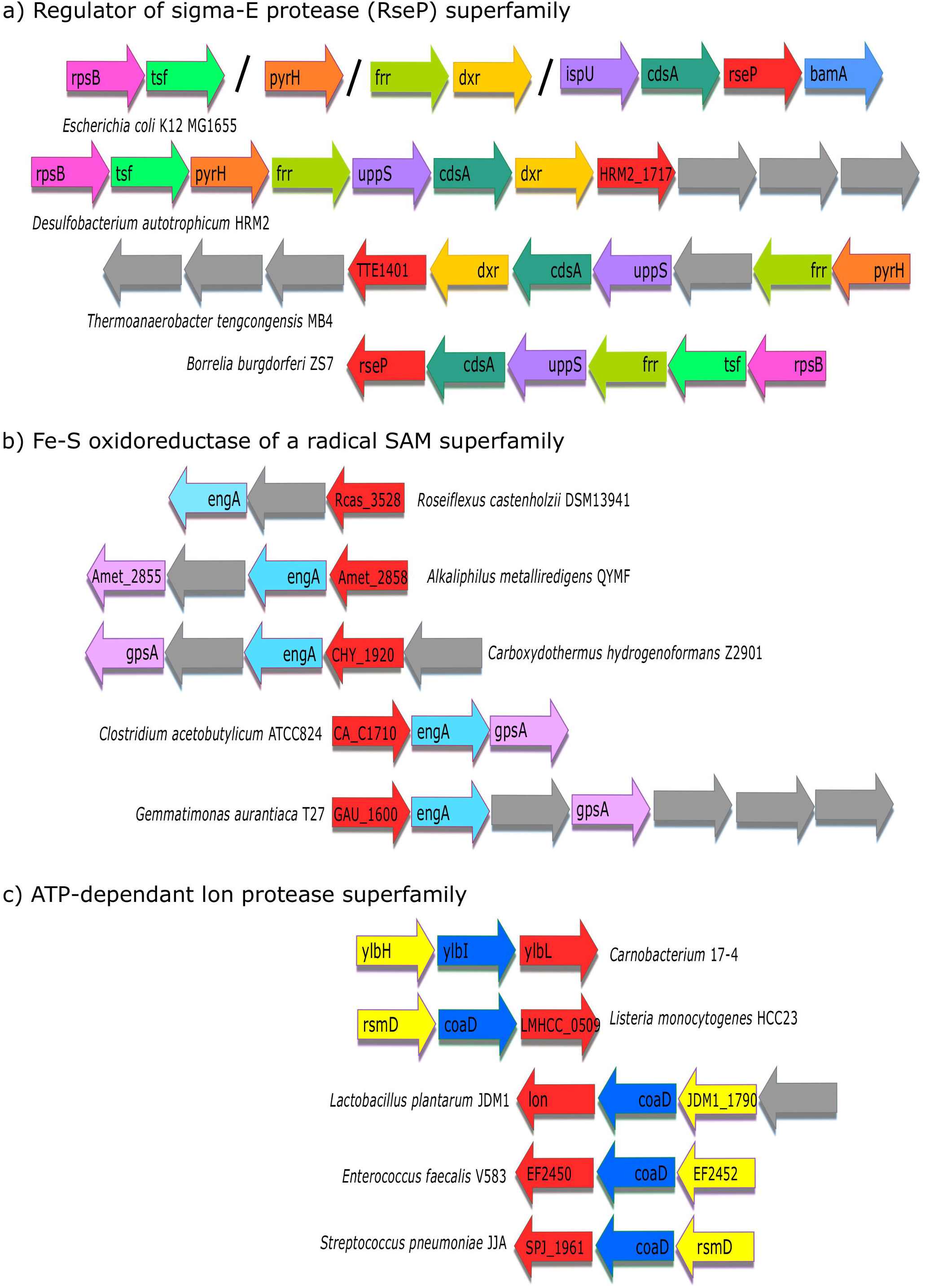
Genomic context reveals co-localization of PDZ encoding genes with translation-associated genes. a) RseP family members are frequently associated with cdsA, uppS (ispU), dxr, frr, rpsB and bamA genes. Intergenic distance greater than 50 nucleotides is indicated by slash (/). b) Fe-S oxidoreductases of Radical SAM family members occur in the vicinity of engA (der) and gpsA genes. The gene between engA and der is conserved in many species but no function is defined for it. c) ATP-dependent lon protease family members are well conserved with coaD and rsmD gene. The rsmD gene is not well annotated in many genomes. Proteins encoded by dxr, rpsB, frr, der, and rsmD are associated with translation related functions. The genes are included in this analysis if they are placed within a distance of 50 nucleotides from the PDZ-coding gene with the exception of *Escherichia coli* where it was 300 bases (a). Locus identifiers were used when gene names were not available. Red color is used to represent PDZ-domain containing proteins in each panel. Homologous genes are represented in the same color. Grey colored genes are not conserved in the neighborhood. Arrows pointing towards the right represent the plus strand and towards the left represent the minus strand of the genome.

The genomic context of Fe-oxidoreductase family members contains gpsA (encodes a glycerol-3-phosphate dehydrogenase enzyme), and engA (Fig. 6b). The engA gene encodes a GTPase also called Der, required for ribosome assembly and stability. It co-transcribes with the outer membrane protein BamB coding gene from a σ24 promoter in *Escherichia coli* (Rhodius et al. 2006). The product of bamB is part of the large multi-protein BAM complex responsible for outer membrane biogenesis, including bamA sharing genomic context with rseP (Fig. 6a). Interestingly, the radical SAM domain predicted to have three-dimensional fold similar to SAM methyltransferase involved in translation. The genomic neighbourhood of this family with Der further strengthen our hypothesis that the radical SAM family is likely to be involved in translation related functions.

PDZ domain is combined with ATP-dependent lon protease in 292 uncharacterized proteins of actinobacteria and firmicutes (Table 1). The prototype example of such domain organization is the *Bacillus subtilis* membrane protein YlbL which has a structural fold similar to ribosomal protein S5. The genes encoding these proteins share genomic neighbourhood with rsmD (Fig. 6c). The rsmD product plays a critical role in the methylation of G966 of 16S rRNA (Weitzmann et al. 1991), which modulates the early stages of translation initiation (Burakovsky et al. 2012). Substitutions at G966 generally have no effects on cell growth (Jemiolo et al. 1991) but on stress response (Burakovsky et al. 2012). It has been suggested that RsmD acts in a complex way to shape the bacterial proteome in stress conditions. The null mutants of rsmD gene remain in lag phase for prolonged periods under stress (Burakovsky et al. 2012; Christensen et al. 2001). Interestingly, this complements the inhibitory effect of lon proteases on translation, particularly in nutritional stresses (Christensen et al. 2001).

## Discussion

The present study proposes a concrete classification of 97% PDZ proteins into twelve families. Of these, six families have at least one characterized protein, while remaining six are uncharacterized till date. A combination of PDZ and protease domains is a common theme observed in 88% of classified proteins. PDZ-protease domain combination seems to have appeared later in the evolution. Prior to this event the function of PDZ domains was presumably to bind iron available in large quantity in early earth atmosphere (Fig. 5e). The gain of peptide recognition ability of PDZ domains is likely to have facilitated their evolution along with a variety of proteases, suggesting their major role in providing substrate-specificity towards controlled proteolysis. HtrA protease family being present in almost all investigated genomes (except few cell wall-less bacteria) while previously reported in eukaryotes including plants, and animals (Clausen et al. 2011; Ponting and Phillips 1995; Ponting 1997; Sakarya et al. 2010; Schuhmann et al. 2012), suggests cellular heat response is universally conserved. HtrA proteases first diverged from non-protease PDZ domain-containing proteins, and shared last common ancestor with remaining protease families (Fig. 5e). This is consistent with their heat resistance and requirement in higher temperature of early earth. The DUF3340 domain of Ctp protease family proteins is homologous to many metazoan IRBPs suggesting link between them. Furthermore, PDZ domains of many Ctp family proteins were predicted to be canonical/metazoan form. This is consistent with their recent divergences evident from the phylogenetic analysis (Fig. 5e). Proteins harbouring Aminopeptidase N, ATP-dependent lon protease, Aspartyl protease, and Zn-dependent exopeptidase domains are yet to be functionally characterized, along with non-proteases Fe-S oxidoreductases and Haem-binding uptake, Tiki superfamily.

In search of their putative functions, we analyzed genomic neighbourhood all protein-coding genes. Conserved neighbourhood were observed for three families, of which the ATP-dependent lon proteases (structural similarity with tRNA or rRNA modification domain) and Fe-S oxidoreductases are often placed with genes coding for proteins involved in translation and membrane biogenesis, indicating their functions in these processes. RseP intra-membrane metalloproteases are already well characterized at experimental levels, and are also placed with genes involved in above-mentioned functions. Such consistent chromosomal proximity of PDZ domain coding genes reinforces their roles in translation and membrane biogenesis. If PDZ domains are indeed involved in translation and membrane biogenesis, the coupling of translation and transcription in prokaryotes might explain the depletion of non-canonical PDZ from eukaryotes wherein these processes are spatio-temporally isolated and such domains might not have provided functional advantages. Membranes are the first line of defence for eubacteria against fluctuations in the extracellular milieu. PDZ domain-containing proteins sense unfolded proteins during various stresses and activate specialized σ24 response. It would be worthwhile to experimentally explore how PDZ proteins modulate translation efficiency and remodel membranes during stress in addition to their probable role in tRNA and rRNA modifications.

The expansion of PDZ domains in planctomycetes (debatable host in endosymbiosis theory of first eukaryotic cell formation) and myxococcus having many metazoan/eukaryote features supports the hypothesis that PDZ domains might have co-evolved with multi-cellularity/complexity (Harris and Lim 2001). However, the origin of PDZ domains was unclear in metazoa as well as in prokaryotes. We hypothesized the PDZ domain of the Fe-S Oxidoreductase family as the probable ancestor for all PDZ domains that might have helped the ancestral eubacterial species to withstand the anaerobic atmosphere of early Earth. Their radical SAM domains might have provided the means of reductive energy generation or translation stability under anaerobic atmosphere of early Earth. Oxygen availability might have negatively selected these proteins in the species diverged from ancestral gladobacteria but retained them in some extant obligatory anaerobes. One the other hand, proteases might have expanded with the availability of oxygen and help in adapting terrestrial niche (Fig. 3).

Though we provide evidence of ancestral relationship between bacterial and metazoan PDZ domains as well as presence of hundreds of canonical PDZ domains (in Ctp protease family) in eubacteria, their depletion or absence in archaea and fungi hinders solid explanation for this transition. Based on the analysis presented here we argue that Archaea presumably never had canonical PDZ domains. The bacterial endosymbiont might have contributed these domains to eukaryotic phylogeny, where they were lost recurrently only in fungi and ecdysozoa. Phylogenetic analysis confirms recent divergence of the PDZ domains of Ctp family, which might have contributed canonical PDZ form on metazoan phylogeny. Collectively, our comprehensive analysis provides insights into the emergence of the PDZ domains and their functional divergence during evolution.

## Methods

### Identification and analysis of PDZ domain-containing proteins in completely sequenced genomes

Protein sequence and annotation files were obtained from the National Centre for Biotechnology Information (NCBI) for completely sequenced prokaryotic and fungal species (Sayers et al. 2012). Out of total 2,057 genomes we selected 1,474 representative genomes for which phenotypic information was available for further analyses. Genome size, taxonomy and phenotypic information for these species were obtained from ftp://ftp.ncbi.nih.gov/genomes/genomeprj/. Hidden Markov Models (HMM) of PDZ domains were downloaded from Pfam and Superfamily databases (Gough et al. 2001; Sonnhammer et al. 1997). The accession numbers for Pfam domains are PF00595, PF13180, PF12812 and 50156 for Superfamily database. These HMMs were searched against all protein sequences of each selected genome using *hmmsearch* program from HMMER package (Finn et al. 2011). The inclusion thresholds of e-value <= 0.01 and 0.03 were used to consider the significance of output sequence and obtained hit respectively. The resulting sequences were subjected to the *hmmscan* analysis to identify other domains using HMMs of complete Pfam and Superfamily databases. The e-value was set at 0.01 for HMM and 0.01 for obtained hits. The output of *hmmscan* for both Pfam and Superfamily database search was analyzed separately using Perl scripts to extract the domain organization of each protein sequence. This search identified 7,852 protein sequences with at least one PDZ domain predicted either using Pfam or Superfamily HMM model. Sub-cellular localization of these proteins was predicted using Phobius web server (Kall et al. 2007). The data was processed using in-house Perl scripts and visualized over pruned version of NCBI taxonomy tree which was created using interactive Tree Of Life (iTOL) web server’s API tool by providing NCBI taxonomy identifiers for investigated organisms (Letunic and Bork 2007).

### Classification of PDZ domain-containing proteins

The classification of PDZ domain-containing proteins is challenging, owing to their sequence and structural variations. On several instances we were unable to find correspondence between hits identified by Superfamily and Pfam models due to the different classification strategies adapted by them. To overcome this problem, Pfam classification was used as a reference and always cross-checked with Superfamily classification for consistency. First, we grouped proteins based on conserved Pfam domain architectures using in-house Perl scripts. The remaining sequences were manually checked and assigned to each group. Second, the *clustalo* program was used to construct a multiple sequence alignment with default settings for each group which was manually analyzed to exclude highly divergent sequences (Sievers et al. 2011). At multiple instances a prototype motif of specific family was considered for assigning proteins to their respective group (Supplementary Fig. 6 & 7). This semi-automatic sequence analysis led to classification of 7,318 proteins out of 7,852 into 12 families. We were unable to classify 3% proteins due to their presence in less than 20 species and highly variable domain combinations.

### Sequence, Structure and Phylogenetic analysis

PDZ domains are inherently diverged at sequence and structure level. This hinders the phylogenetic signal in addition to its small length leaving few informative sites for phylogenetic reconstruction. Therefore, we selected PDZ domains only from delta-proteobacteria group to reconstruct phylogeny. The selection was based on the presence of both GspC and radical SAM family proteins in them. *clustalo* program was used to align sequences using Superfamily HMM model, which is based on the alignment of PDZ domain structures. Positions that were conserved in more than 70% sequences were retained for analysis. We manually monitored MSA to remove sequences that were highly diverged. Phylogenetic trees were reconstructed with Fitch-Margoliash and parsimony algorithms available through *fitch* and *protpars* programs in Phylip package respectively (Felsenstein 1989). The statistical significance was accessed with 1,000 bootstraps. Protein distance matrix was constructed using *protdist* program in Phylip package to feed in *fitch* program. A maximum likelihood tree was constructed using RAxML v. 8.1.24 (Stamatakis 2006), as implemented on the CIPRES web server (Miller 2010), under the WAG plus gamma model of evolution, and with the number of bootstraps automatically determined (MRE-based bootstrapping criterion). A total of 660 bootstrap replicates were conducted under the rapid bootstrapping algorithm, with 100 sampled to generate proportional support values. MrBayes analysis was performed for one million generations with WAG substitution model and gamma distribution for four categories (Huelsenbeck and Ronquist 2001). The trees were sampled after every 1000 generations and the first 25% was discarded as burn-in. The resulting trees were visualized in FigTree1.4 (available from http://tree.bio.ed.ac.uk/software/figtree/) with levels of support shown as posterior probabilities. The structure was predicted using Phyre2 web server (Kelley et al. 2015) and Pymol software was used for analysis and visualization.

### Genomic context analysis

Protein annotation files were used to extract genomic coordinates of genes encoding PDZ-domain containing proteins. We considered a gene as a neighbor, if it is co-directional and placed within 50 nucleotide bases from a PDZ domain-coding gene on the genome (Muley and Ranjan 2012).

### Statistical significance

The difference between two distributions of PDZ containing proteins/domains was assessed using Wilcox rank sum test (Wilcoxon 1945). Null hypothesis was either no difference in two distributions or one distribution is greater than the other. The null hypothesis was rejected and the difference considered significant if the Wilcox test p-value was lower than 0.05.

## Acknowledgements

The authors wish to thank Anne Hahn for help in editing the manuscript and Amol Kolte for help in preparing Figure 4. This work was supported by the Center of Excellence in Epigenetics grant [BT/01/COE/09/07] from Department of Biotechnology, India; Indian Institute of Science Education and Research, Pune, India [core grant] (to SG). VYM was supported by the Swarnajayanti fellowship to SG.

## Authors’ Contributions

VYM conceived the study. VYM designed and performed analysis. VYM interpreted results. SG supervised the study and contributed to the interpretations. VYM and SG wrote the manuscript.

